# The putative phosphate transporter PitB (PP1373) is involved in tellurite uptake in *Pseudomonas putida* KT2440

**DOI:** 10.1101/2020.08.24.265637

**Authors:** Rafael Montenegro, Sofía Vieto, Daniela Wicki-Emmenegger, Felipe Vásquez-Castro, Carolina Coronado-Ruiz, Paola Fuentes-Schweizer, Paula Calderón, Reinaldo Pereira, Max Chavarría

## Abstract

Tellurium oxyanions are chemical species with great toxicity; their presence in the environment has increased because of mining industries and photovoltaic and electronic waste. Recovery strategies based on microorganisms for this metalloid are of interest, but further studies of the transport systems and enzymes responsible for implementing tellurium transformations are required because many mechanisms remain unknown. Here, we investigated the involvement in tellurite uptake of the putative phosphate transporter PitB (PP1373) in soil bacterium *Pseudomonas putida* KT2440. For this purpose, through a method based on the CRISPR/Cas9 system, we generated a strain deficient in *pitB* gene and characterized its phenotype on exposing it to varied concentrations of tellurite. Growth curves and Transmission Electronic Microscopy experiments of wild type and Δ*pitB* showed that both strains were able to internalize tellurite into the cytoplasm and reduce the oxyanion to black nano-sized and rod-shaped tellurium particles, however, Δ*pitB* strain showed an increased resistance to the tellurite toxic effects. At a concentration of 100 uM tellurite, where the biomass formation of wild type strain decreased by half, we observed a greater ability of Δ*pitB* to reduce this oxyanion with respect to wild type strain (~38% vs ~16%), which is related by the greater biomass production of Δ*pitB* and not by a greater consumption of tellurite per cell. The phenotype of the mutant was restored on over-expressing *pitB* in trans. In summary, our results indicate that PitB is one of several transporters responsible for tellurite uptake in *P. putida* KT2440.

## Introduction

Tellurium oxyanions, namely tellurate (TeO_4_^2−^) and tellurite (TeO_3_^2−^), are chemical compounds with great toxicity, which are found naturally in small concentrations, but its presence in the environment has increased because of mining industries and photovoltaic and electronic waste [1,2]. Interest has hence increased in the development of bioremediation and recovery strategies for tellurium as well as for other metals and metalloids. In turn, the global demand for this element in technology manufacture, drives a search for new environmentally compatible tools for the recovery of this element from its natural sources [3].

Despite the toxicity of these oxyanions (tellurite is considered one of the most toxic metalloid oxyanion [2,4,5]), many environmental microorganisms have shown a remarkable ability to resist and to transform tellurium compounds [2,4,6]; which hints that microbial cells might become a biotechnological tool for the recovery of this element. The toxic effect of TeO_3_^2−^ in bacteria occurs at concentrations as small as 1 μg mL^−1^ (4 μM) for *Escherichia coli* [4,5], or 2500 μg mL^−1^ (10 mM) for resistant strains such as *Erythromicrobium hydrolyticum* [7].

Two biotransformations of tellurite have been reported in bacteria: (i) the reduction of tellurite to elemental tellurium (forming black nanoparticles) and (ii) the formation of gaseous compound dimethyl telluride ((CH_3_)_2_Te) (2,4,8,9). The formation of tellurium nanoparticles was first reported in *Rhodobacter sphaeroides* by Moore & Kaplan in 1992 [4,10]; it has since been described in several bacteria with varied sizes and shapes [4,8,11–14]. Although there exists some knowledge about genes, enzymes and metabolic steps involved in the metabolism of tellurite, the process is poorly understood in most microorganisms. Tellurite is known to activate the machinery of resistance to oxidative stress in *E. coli* and *Pseudomonas pseudoalcaligenes* that includes NADPH/glutathione-mediated reactions [2,4,13,15]. In *Rhodobacter capsulatus,* a thiol:disulfide oxidoreductase was shown to participate in electron transport resulting in the reduction of tellurite [16]. Several tellurium-resistant genes have been identified in *E. coli* (*ter, kilA and tehAB operons)* [5] and other bacteria such as *R. sphaeroides (trgAB genes* and *nap* gene), *Pseudomonas syringae (tmp* gene*), Lactococcus lactis (trmA* gene*), Geobacillus stearothermophilus (ubiE* gene*), Proteus mirabilis (terC* gene), and *Alcaligenes spp.* (*ter* genes) [2,4,17].

One aspect not defined for all bacterial systems regarding the tellurite metabolism is its transport system. Some transporters have been identified to have tellurite unspecific activity, such as the low-affinity inorganic phosphate transporter (PitA) in *E. coli*, the acetate permease (ActP) in the facultative phototroph *Rhodobacter capsulatus* [4,18] and the PstA and PstD proteins (involved in phosphate transport) in Gram-positive bacterium *Lactococcus lactis*. Specifically, in *E. coli* the low-affinity inorganic phosphate transporter (PitA) was demonstrated to participate in the tellurite uptake of this bacterium, whereas its homologous (81 % identity) PitB is not involved [19]. The authors of that work reported that *pitA* mutation increases four-fold the tolerance to tellurite and that the intracellular uptake was decreased by half that of the wild type. Elías et al (19), deleted the *pitB* gene, but the mutant exhibited a behavior similar to that of the wild type, in terms of uptake and sensitivity to tellurite. In *R. capsulatus* the acetate transport system ActP was demonstrated to be utilized for tellurite uptake and that a limitation of its transport (by mutation or adding acetate in the culture medium) greatly increased the resistance to the oxyanion (20). In the case of *E. coli*, the ActP transporter showed tellurite uptake activity, but mainly during early exposure, whereas PitAB remained active in longer exposures [21]; this fact indicates that the main uptake is always mediated by the PitA transporter.

*Pseudomonas putida* KT2440 is a model microorganism, showing metabolic advantages for the bioconversion and bioremediation of various chemical compounds, such as aromatic compounds (toluene, m-xylene), and metalloid ions among others [22–25]. *P. putida* KT2440 has a specialized metabolism (i.e. EDEMP cycle) that generates an increased capacity to produce more equivalents of reducing power (i.e. NADPH), thus counteract oxidative stress [26–28]. This characteristic makes it an ideal microorganism to perform biotransformations that require reducing power, such as the reduction of tellurite and its recovery as elemental tellurium. A previous report demonstrated that *P. putida* KT2440 transformed with plasmids containing tellurite resistance genes (Tel^R^; specifically *kilA* and *telAB*) was able to grow and form elemental tellurium at concentrations between 24-315 μM (6-80 μg mL^−1^) [29]. Other members of genus *Pseudomonas* can act efficiently as microbial systems that remove tellurite (9,30). Taking these facts into account, we consider relevant an exploration of the use of bacterium *Pseudomonas putida* as a microbial system for the biotransformation of tellurite to tellurium. Such an objective implies studying the wild-type intrinsic metabolic and transport machinery to evaluate whether it can be improved through metabolic engineering strategies. In this work, we studied the function of the low-affinity inorganic phosphate (Pit) transporters of *P. putida* KT2440 for tellurite uptake. The *P. putida* KT2440 genome [25] reveals the existence of a single Pit transporter (PitB encoded by PP1373 gene) unlike *E. coli* in which one can find two systems (PitA, PitB). For this purpose, we generated a strain deficient in the PP1373 gene, encoding the putative phosphate transporter PitB, through a method based on the CRISPR/Cas9 system [31]. We evaluated the function of the product of the PP1373 gene through growth curves, Transmission Electronic Microscopy (TEM) experiments and the kinetics of tellurite uptake. A plasmid overexpressing the *pitB* gene was constructed to perform complementary studies.

## Materials and Methods

### Bacterial strains, culture media and growth conditions

*P. putida* KT2440 and the mutant strain Δ*pitB* were grown aerobically at 30 °C with orbital shaking at 180 rpm either in LB broth [32] or in M9 medium containing citrate (0.2% w/v) [32] as the sole source of carbon and energy. Cultures in plates were prepared using the same media supplemented with agar (1.6% w/v). When required, potassium tellurite (Sigma-Aldrich WGK, Germany) was added at concentration 10 μM – 1000 μM.

### Construction of *P. putida* Δ*pitB* strain

Guided mutagenesis was accomplished with a CRISPR/Cas9 recombining tool previously described [31], using a three-plasmid strategy: pSEVA421-Cas9tr, pSEVA658-ssr and pSEVA231-CRISPR. A guide sequence of 30 nucleotides (5’-AAACTGGTATAGATGACGGTAGCTACCGCGTTGGG-3’) was searched in gene PP1373 containing a PAM motif that was synthesized in sense and anti-sense sequence; after annealing the fragment, it was cloned into plasmid pSEVA231-CRISPR with BsaI restriction and ligation. An oligo recombining 90 nucleotide bases was synthesized (5’-TATGAAAAAGGCGCCCGAGGGCGCCTTCATTCATTGCGGTGTCACGAAGGTTGTCTGGCGGGTCTTAGGGGGGCGGGAATTATGCCAGAAA-3’) containing sequences upstream and downstream of the PP1373 gene, so as to eliminate the *pitB* (PP1373) gene as part of a recombination event. All mutagenesis was performed as was described by Aparicio et al. [31]. Mutagenesis was verified with PCR of primers 1373-check-F 5’-GATGGGGTCGTTGTTAGGGG-3’ and 1373-check-R 5’-TTCAGCCAGGAACTGGAACGC-3’, which flanked the whole *pitB* gene.

### Cloning of *pitB* gene from *P. putida* KT2440 for complementation

For the complementation of the mutant, the DNA sequence encoding the *pitB* gene (PP1373) of *P. putida* KT2440 was amplified directly from the chromosome of the bacterium with primers 1373-F-(5’-TGCCGGAATTCATGATCGATTTATTCAGCGGACTG-3’ and 1373-R 5’-AACCCAAGCTTTCAGCCAATGGCTTTGGACG-3’). The resulting amplicon was digested, ligated to the polylinker of the expression plasmid pSEVA438 (33) as an EcoRI HindIII fragment (producing pRMM-*pitB*) and subsequently transformed and propagated in *E. coli* DH5α. To verify the ligation, we replated colonies in fresh media with Streptomycin (Sm 100 μg mL^−1^); a colony PCR was performed with the 1373-F and 1373-R primers. After verifying the ligation of the plasmid pSEVA438 with the *pitB* gene, we transformed the construct into strain *P. putida* Δ*pitB*, whereas the empty backbone was transformed to both strains *P. putida* KT2440 and Δ*pitB* for use as a control in the experiments.

### Bacterial growth in presence of tellurite

To establish suitable conditions to analyze the sensitivity to tellurite we grew preinocula of both *P. putida* KT2440 and Δ*pitB* strain overnight in M9 as described above. These cultures were diluted to an initial optical density approximately 0.05 at 600 nm in fresh M9 medium containing a gradient of potassium tellurite (0, 10, 15, 20, 25, 50, 75, 100, 200, 500 and 1000 μM). Bacterial growth was estimated on monitoring the optical density at 600 nm (Synergy H1 Hybrid Multi-Mode Reader, Biotek, Winooski VT, USA) over three biological replicates of cultures in plates (96 wells; Nunclon™ Δ Surface; Nunc A/S, Roskilde, Denmark). Plates were incubated at 30 °C for 24 h with continuous orbital shaking at 180 rpm; the optical density was measured every 10 min. For plasmid-based complementation experiments in a 96-well plate, tellurite was fixed at 100 μM; the media was supplemented as well with Sm (100 μg mL^−1^) and 3-methylbenzoate (3MB, 0.5 mM), added to induce genetic expression.

### Bacterial tellurite sensitivity

To determine the toxicity of tellurite to *P. putida* KT2440 and Δ*pitB* strains, colony-forming units (CFU) were calculated in 25 h cultures of M9 media with tellurite 100 μM for both strains. Cultures were inoculated at 0.05 OD_600_ and incubated at 30 °C for 25 h with continuous orbital shaking at 180 rpm. Samples were taken each 2.5 h; then each sample was serial diluted from 10° to 10^−8^ in Phosphate-Buffered Saline (PBS), plated into M9 agar plates and after 24 h of incubation, colonies were counted and the exact CFU number was calculated. CFU was reported over three biological replicates of the experiment.

### Transmission electronic microscopy energy dispersive X-ray (TEM-EDX) experiments

*P. putida* KT2440 and Δ*pitB* strains were grown in M9 media, both sole and supplemented with tellurite 100 μM, in order to obtain TEM samples. Cultures inoculated at 0.05 OD_600_ were incubated at 30 °C for 25 h with continuous orbital shaking at 180 rpm. Samples were taken at 2.5, 12.5 and 25 hours of culture and in duplicates, based on two biological replicates. Each sample was then centrifuged to remove the media, next the pellet was resuspended in 100 μL of Karnovsky’s fixative. Then, a 5 μL aliquot was taken and placed on a copper grid with formvar and carbon coated and let dry for 5 minutes. Images and X-ray microanalysis were performed using HT-7700 (Hitachi, Japan) and JEOL JEM 2011 (JEOL, Japan) transmission electron microscopes (TEM).

### Determination of tellurite uptake by inductively-coupled-plasma optical-emission spectrometry (ICP-OES)

*P. putida* KT2440 and the mutant strain cells were cultured in conical flasks (3 L) filled with M9 medium containing citrate (0.2 % w/v) as the sole source of carbon and energy (500 mL) and supplemented with potassium tellurite (100 μM) at 30 °C and 180 rpm for 15 h. To have replicates of each sample the experiment was performed three times. Cell suspension aliquots (15 mL) were withdrawn at intervals 2.5 h during the bacterial growth for measurement of their optical density at 600 nm in 96-well plates (Synergy H1 Hybrid Multi-Mode Reader, Biotek, Winooski VT, USA). Cells in these aliquots were removed on centrifugation (10 min, 4000 g, 4 °C); the resulting supernatants were transferred to a new tube (15 mL) and stored at −25 °C until further analysis. For the analytical measurements, supernatants were filtered in Syringe Filters (Minisart® RC15 Regenerated Cellulose, Sartorius^®^, 0.45 μm), and analyzed with an inductively-coupled-plasma optical-emission spectrometer (ICP-OES), (Optima 3300-XL, Perkin-Elmer, Norwalk, CT, USA). The equipment was operated in an axial configuration; the wavelength used for tellurium was 214.201 nm. The analyte solutions were prepared from potassium tellurite (Sigma Aldrich, Cat 60539-10G, Lot # BCBW5001, ≥ 95.0 %). Appropriate dilutions were made for the preparation of the standards (10-120 μmol L^−1^), which were stored in polyethene flasks under refrigeration. Quality-control measurements were made for blank solution and control solution every fifteen measurements.

## Results and discussion

The genome of the soil bacterium *Pseudomonas putida* [25] bears an ORF for a protein (PP1373) of 490 amino-acid residues (coordinates 1563856-1565328; Fig. 1) and 52.4 KDa MW that shares 55 % identity with the *E. coli* PitA protein and 54 % identity with the *E. coli* PitB protein (Fig. S1). All these criteria qualify PP1373 as a bona fide orthologue of the two enterobacterial counterparts (i.e. both PitA and PitB proteins from *E. coli*). In other *Pseudomonas* species it is possible to find an orthologue annotated as *pitA* with large percentages of identity with *P. putida pitB* (Fig. S2), but, to the best of our knowledge, there is no study with biochemical validation of these transporters in *Pseudomonas* species. Phylogenetic analysis of *pit* genes (Fig. S3) indicated that the two versions of Pit in *E. coli* are the product of a duplication event, whereby they have disparate functions. This condition is consistent with the previously observed differences between both proteins in tellurite transport; whereas PitA seems to be involved in tellurite transport, PitB seems not [19]. *pitB* has been described as having a regulation different from that of the *pitA* gene in *E. coli*, as the former is induced by large levels of inorganic phosphate (Pi) whereas the *pitA* gene is induced by P_i_ starvation and intracellular Zn (II) levels, as it co-transport metals ions; both transporters share the same function of P_i_ transport [34].

**Figure 1.**
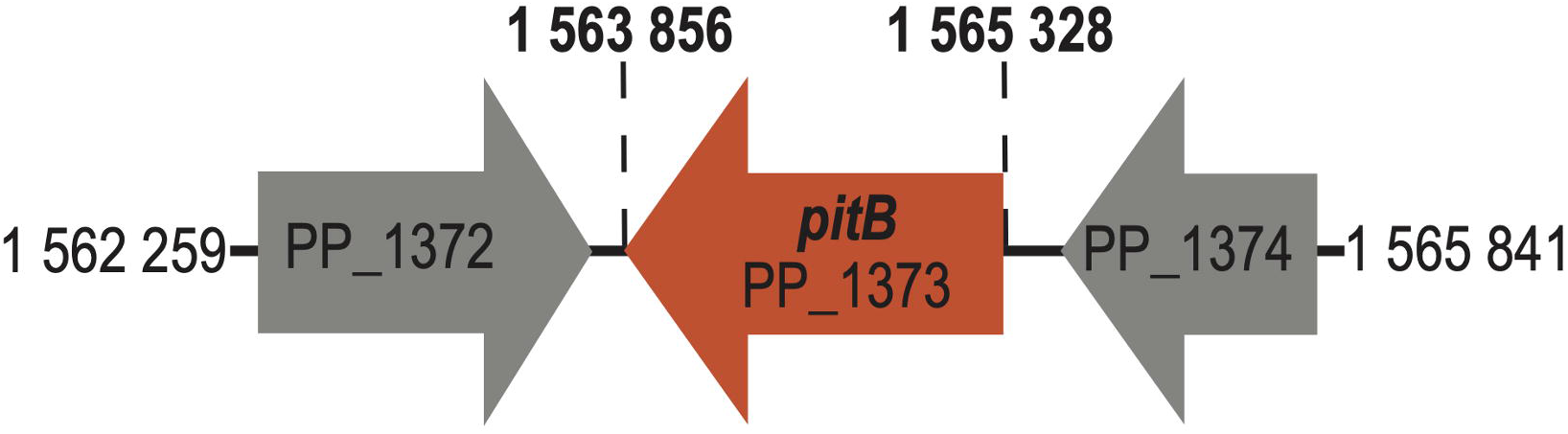
Genomic context of the *pitB* gene (PP1373) in *Pseudomonas putida* KT2440 genome. The genome details of the gene context were retrieved from the *Pseudomonas* Genome Data Base webpage that showed the presence of a single low-affinity inorganic phosphate transporter (*pit*) gene in the KT2440 strain genome.

Despite that the main reported function of *pit* genes is phosphate transport, our interest was focused on its role in the tellurite uptake, as explained above. For this purpose, we constructed a strain deficient in *pitB* gene through a method based on CRISPR-Cas9 and measured the differences in tellurite resistance with respect to the wild type strain. Sensitivity tests (Figs. 2A and S4) with concentrations between 0 and 1000 μM of tellurite revealed that at concentration 200 μM the growth of the wild-type strain was highly inhibited. To obtain the same effect in the Δ*pitB* variant, a concentration 500 μM was required. Our results showed that the tellurite concentration to inhibit growth in *P. putida* KT2440 is larger than the necessary concentration in *E. coli* (50 μM) [19] in a minimal liquid medium. The toxic effect of tellurite has been described to be related to the formation of ROS species causing first oxidative stress; eventually it could lead to cellular death [33,34]. The greater resistance of *P. putida* than of *E. coli* is possibly related to the ability of *P. putida* to resist oxidative stress through its specialized central metabolism [26,27,37].

**Figure 2.**
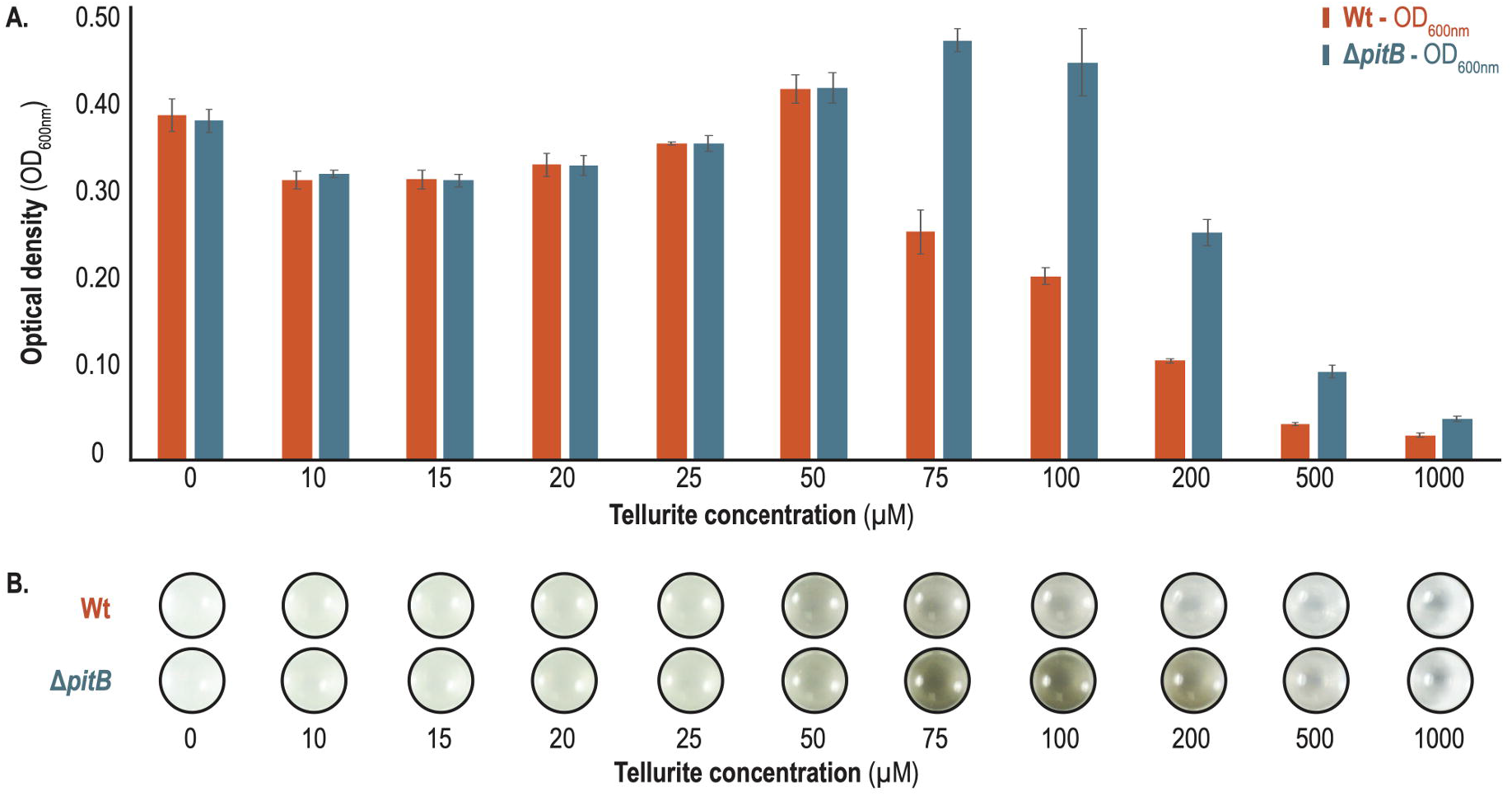
Growth of *P. putida* KT2440 wild-type and Δ*pitB* strains in M9 minimal media with tellurite at varied concentrations. **(A)** Optical Density measurements at 600 nm at 24 h of culture for potassium tellurite at varied concentrations (0 μM to 1000 μM). The results shown correspond to three independent experiments. The phenotypical analysis showed growth inhibition for the wild-type strain beginning at tellurite (75 μM); Δ*pitB* exhibited the same inhibition until reaching 200 μM of the compound. **(B)** Photograph of the microplate experiment after incubation for 24 h to exemplify the phenotypical growth of both strains.

As Figs. 2A and S4 shown, the sensitivity in *P. putida* KT2440 is initially observed at concentration 75 μM. At that concentration, we obtained a growth rate decreased by 5 % (with respect to wild type in absence of tellurite). In the Δ*pitB* strain a similarly decreased growth rate was initially observed at concentration 200 μM. There was direct visual evidence for the Δ*pitB* variant generating elemental tellurium, as the cultures had a black appearance at concentrations 50-500 μM (Fig. 2B). These results are consistent with observations made of other bacteria such as *E. coli*, for which a Δ*pitA* mutant reduced part of the tellurite from the culture medium. This observation hence indicates that PitB is one of several tellurite transport systems in *P. putida*.

At concentrations above 100 μM, the large growth differences between the wild type and the Δ*pitB* strains allowed us to observe a clear phenotype, even visually. Considering the results of Fig. 2, in which a clear phenotype is observed between the wild type and the mutant strain at tellurite 100 μM, we selected this concentration for further experiments. As seen in Fig. 2B, photographs of the microplates after growth for 24 h reveal that, at concentration 100 μM, Δ*pitB* produces more elemental tellurium (evident from a greater blackness of the medium) than the wild-type strain. This result could seem inconsistent if we consider that PitB is a tellurite transporter and that its deletion should decrease the uptake of the oxyanion and thereby generate less elemental tellurium. However, this result makes sense considering the sensitivity of tellurite on both strains at a concentration of 100 μM (Fig. 3A), where the wild type showed an increase toxicity leading to a reduction in CFU number, while the Δ*pitB* strain maintained the resistance at the whole 25 hours of culture. Although, in the wild type strain all tellurite entry systems are functional (including PitB), its poor growth causes small percentages of tellurite conversion; it is hence possible to observe only a slight blackening in liquid experiments (Fig. 2B). Although the Δ*pitB* strain should have a smaller tellurite uptake per cell, it grows much better at 100 μM, reaching optical densities and CFU three times that of the wild type strain. The greater number of cells generates a greater conversion of the tellurite from the medium, increasing the production of elemental tellurium. Experiments in a solid medium (Fig. 3B) are consistent with this explanation. As seen in Fig. 3B, the wild-type strain is barely able to grow forming a few small black colonies, whereas the Δ*pitB* variant has appreciable growth and hence greater rates of tellurite conversion.

**Figure 3.**
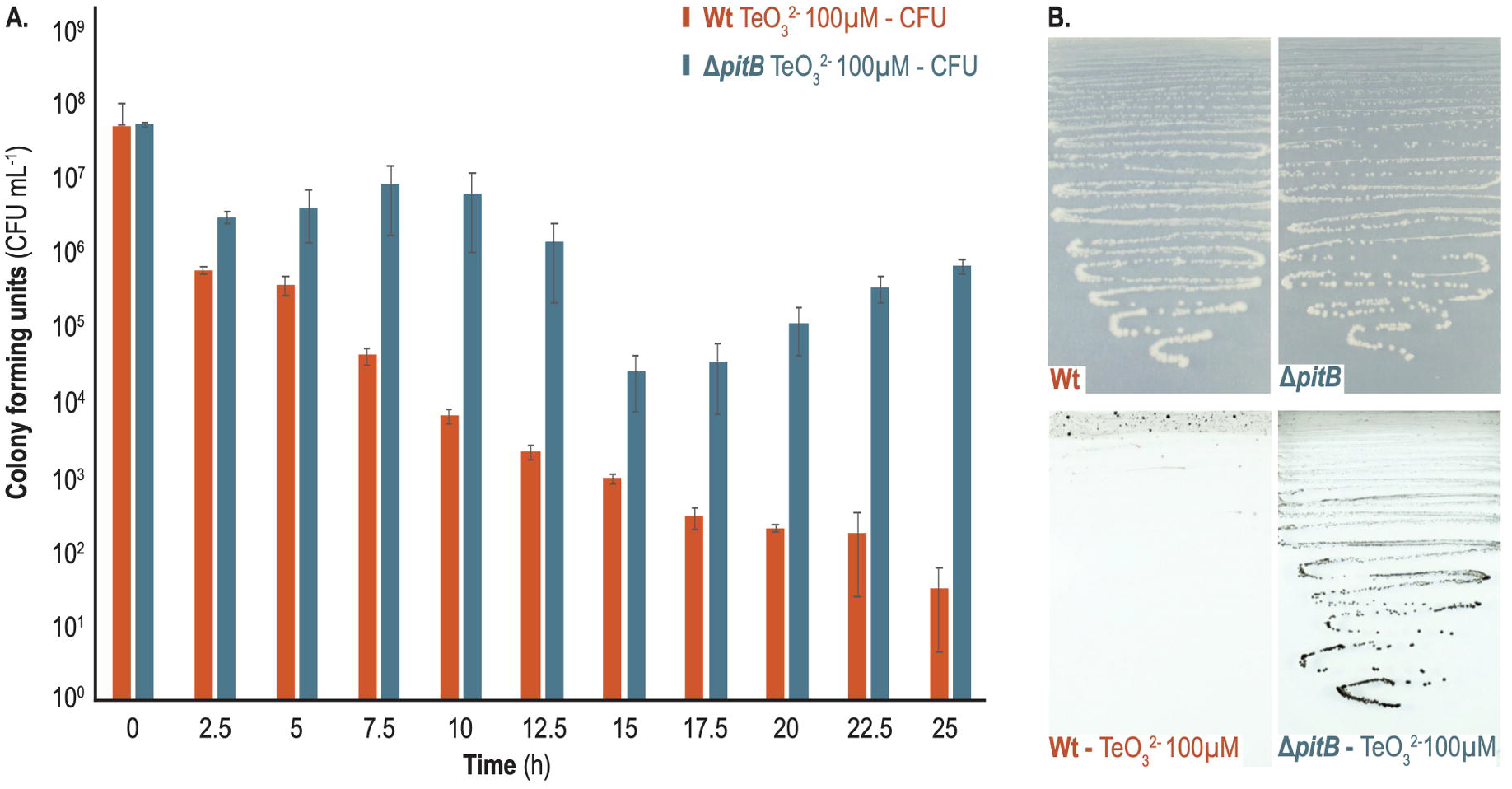
Sensitivity of *P. putida* KT2440 wild-type and Δ*pitB* strains to tellurite 100 μM. **(A)** Toxicity to tellurite was defined by calculating the colony forming units (CFU mL^−1^) on a 25 h culture with tellurite 100 μM, taking samples each 2.5 h for both strains. The wild type strain showed a higher sensitivity to the metalloid decreasing the CFU number constantly over time. While the Δ*pitB* strain was more resistant maintaining the CFU number during the first 12.5 hours and after the 15^th^ hour even showing an increase in the CFU number. **(B)** Photograph of the culture in M9 media agar plates in presence and absence of tellurite (100 μM) for both strains at incubation for 48 h. At tellurite concentration of 100 μM, the Δ*pitB* strain was resistant and capable of growing in agar plates, contrary to the wild type.

TEM experiments were carried out in order to observe the location of the tellurium particles in the cells of *P. putida* as well as the shape and size of these particles. Results in Figures 4A and 4B show that the cells treated with tellurite (both in the wild type and Δ*pitB* strains) present the appearance of electrodense nanostructures with the appearance of rods. These nanostructures are observed mainly within the cell and some of them associated with the outer membrane. The nanostructures observed in *P. putida* are similar to those observed in other species such as *Rhodococcus aetherivorans* (38). Energy dispersive X-ray (EDX) analysis (Fig. 4C) identified the presence of tellurium only in the samples treated with tellurite in addition to other elements associated with the cells. Therefore, the TEM-EDX results suggest that (i) tellurium deposits occur at the cytoplasmic level, (ii) elemental tellurium particles are nano-sized and rod-shaped, and (iii) the presence of elemental tellurium in the cytoplasm of the Δ*pitB* strain allows us to unequivocally conclude the presence of another tellurite transporter in *P. putida*.

**Figure 4.**
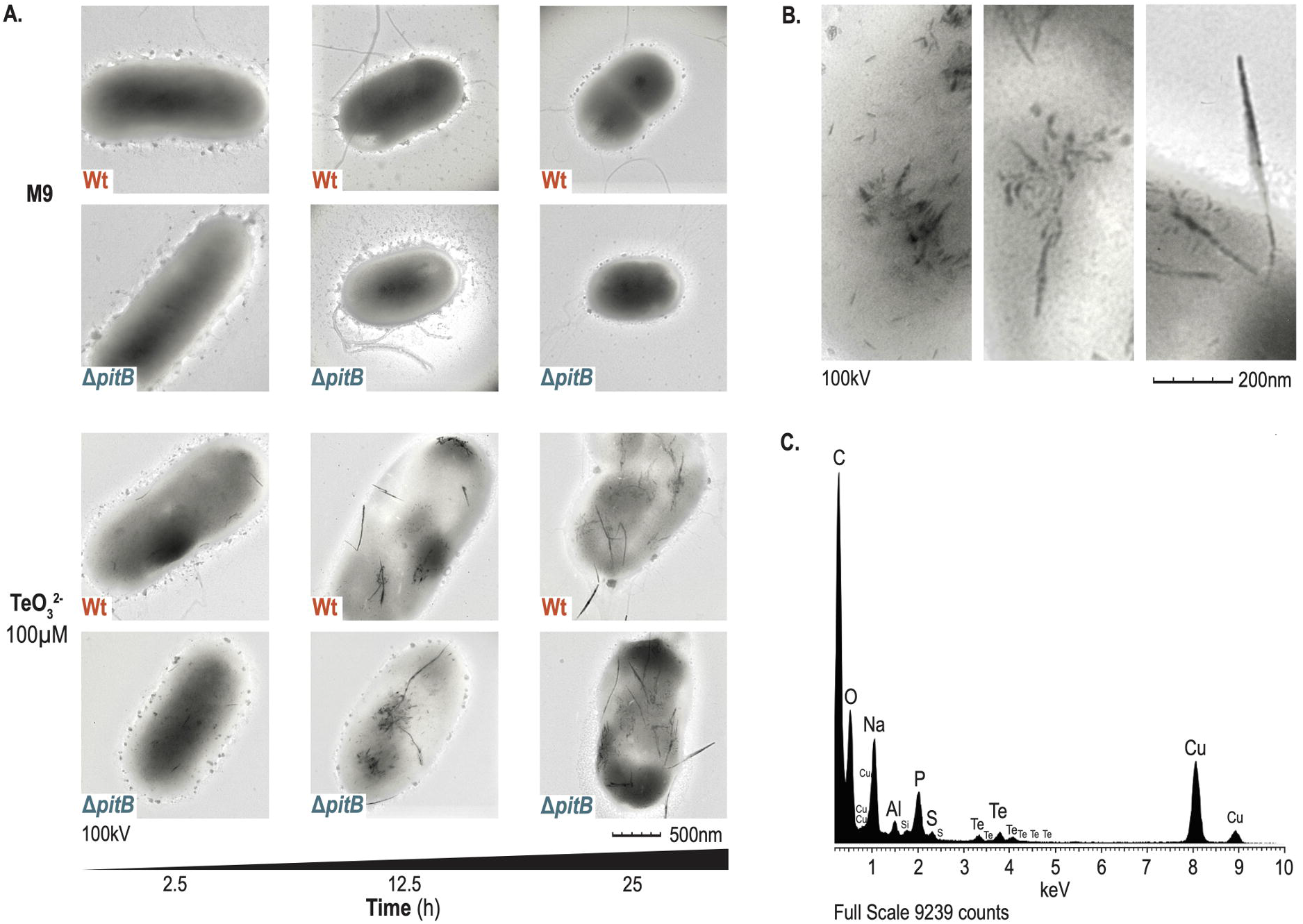
TEM-EDX analysis of *P. putida* KT2440 wild-type and Δ*pitB* strains in sole M9 media and supplemented with tellurite 100 μM. **(A)** Chronological TEM images of KT2440 and Δ*pitB* strains cultured in absence and presence of Tellurite. Images obtained at 100 kV. The visual phenotype was the same for both strains, in M9 media there was not any electrodense structure; while in M9 with tellurite both strains showed electrodense nanostructures inside the cells from 2.5 hour of culture, nanostructures that increase their presence over time. **(B)** TEM close up images of the nanostructures. Nanostructures have sizes ranging from 50 nm to 500 nm. **(C)** EDX analysis revealing the presence of tellurium in the nanostructures.

In order to validate whether the observed differences in growth and formation of tellurium nanostructures were due to the absence of the PitB transporter we run two different experiments: (i) a gene complementation test, and (ii) a comparison of tellurite consumption between the wild type strain and Δ*pitB* mutant. For the complementation test, we constructed plasmid pRMM-*pitB*, in which the *pitB* gene of *P. putida* KT2440 is expressed under the control of a promoter inducible with 3-methylbenzoate (3MB) (see Materials and Methods). This plasmid was introduced into the Δ*pitB* strain; its growth was tested in both solid and liquid media in the presence of tellurite (100 μM). The wild-type strain and the Δ*pitB* variant, carrying the empty vector (pSEVA438), served as controls. As seen in Figure 5A, in the absence of 3MB (*pitB* expression is not induced from the plasmid), the complemented Δ*pitB* strain i.e Δ*pitB* (pRMM-*pitB*) has the same behavior observed for the Δ*pitB* mutant with the empty vector. As expected, when 3MB was added to the culture medium, the expression of the *pitB* gene was induced; the behavior of the Δ*pitB* (pRMM-*pitB*) variant was similar to that of the wild-type strain. The same behavior was observed in the growth curves: when the *pitB* gene was expressed, the sensitivity of the strain to tellurite was restored to the same level as the wild-type strain (Figs. 5B-5C). These results validate what was observed in Figs. 2 and 3 and certify the involvement of the PitB transporter in the tellurite uptake in *P. putida* KT2440.

**Figure 5.**
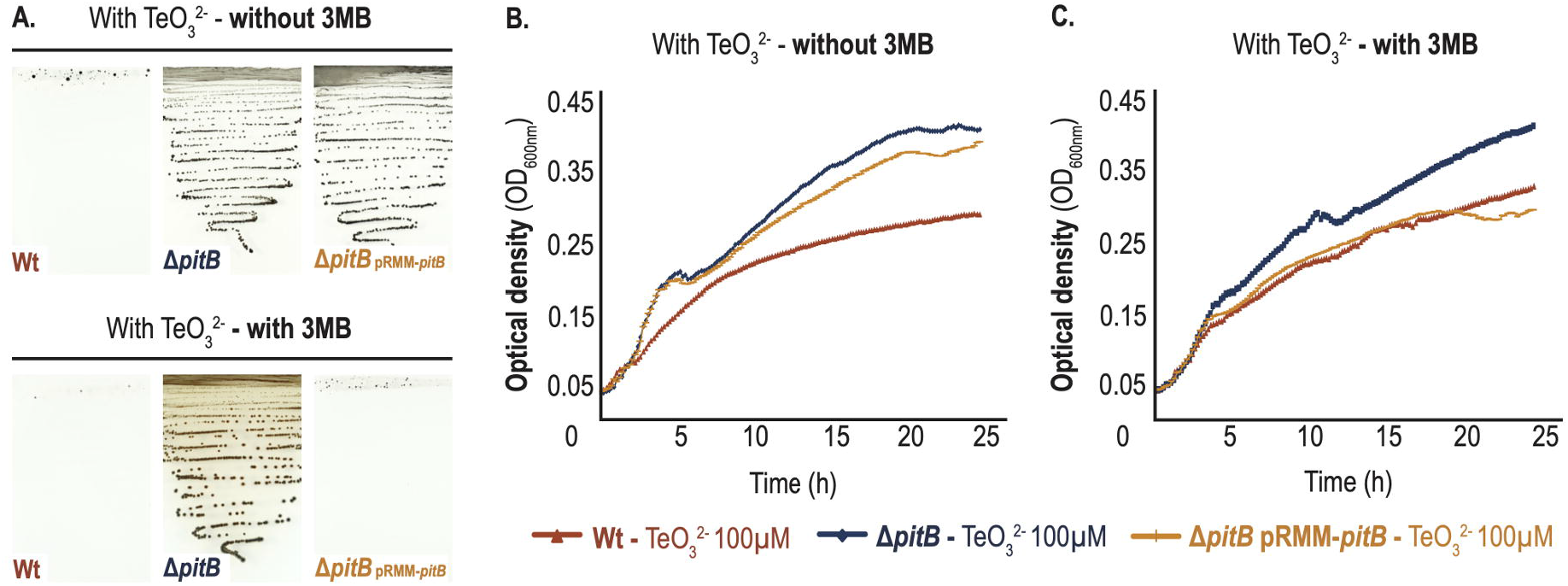
Growth of *P. putida* Δ*pitB* pRMM-*pitB* complemented strain in presence of tellurite. The complementation experiment implied the expression of the *pitB* gene with 3MB as inductor in the Δ*pitB* strain. Strains KT2440 and Δ*pitB* carrying the pSEVA438 vector served as controls. **(A)** Photographs of the culture at 48 h in M9 agar plates with tellurite (100 μM) in absence and presence of 3MB (0.5 mM). **(B)** Growth curves in M9 media with tellurite (100 μM) based on the culture in a 96-well microplate experiment, based on three independent replicates, without 3MB and (**C**) with 3MB (0.5 mM). The curves showed growth inhibition for the KT2440 (pSEVA438) strain with tellurite (100 μM) compared to the Δ*pitB* strains; when 3MB was added to the medium, the Δ*pitB* complemented strain exhibited the same phenotype as the KT2440 (pSEVA438) strain.

For the analysis of the tellurite uptake, we followed the concentration of tellurite (initial concentration ~100 μM) in cultures of *P. putida* KT2440 and the Δ*pitB* strain, during growth for 15 h. For this purpose, we took samples at various times, and then filtered and analyzed with ICP-OES. The results in figure 6 indicate a correlation between growth inhibition and tellurite uptake in *P. putida* KT2440. Before culture for 5 h, the tellurite uptake was greater for the wild type, considering that the depletion of the compound from the media was more rapid, at rate 2.45 μM h^−1^, 18 % greater than for the Δ*pitB* mutant. The KT2440 strain exhibited growth inhibition, for which the optical density measurements remained steady, as well as the tellurite amount remaining in the media. *P. putida* KT2440 (100 μM) was hence able to transform only 16 % of the initial tellurite. The Δ*pitB* strain revealed a different uptake rate, 2.02 μM h^−1^ during the first 5 h of culture and 3.67 μM h^−1^ from the 5 h to the end of the culture; taking tellurite (46.8μM, 38 % of initial concentration), corresponding to 2.3 times that of the KT2440 strain. This observation is consistent with the greatest blackening (tellurium nanostructures) observed for the Δ*pitB* mutant (Figs. 2B and 2C). As discussed above, at concentration 100 μM tellurite is almost lethal for *P. putida* KT2440, forming little biomass; the percentage of conversion of the oxyanion was small (16 %). For its part, at 100 μM, the *pitB* mutant had normal growth and transformed 38 % of the tellurite in the medium, accumulating it as elemental tellurium.

**Figure 6.**
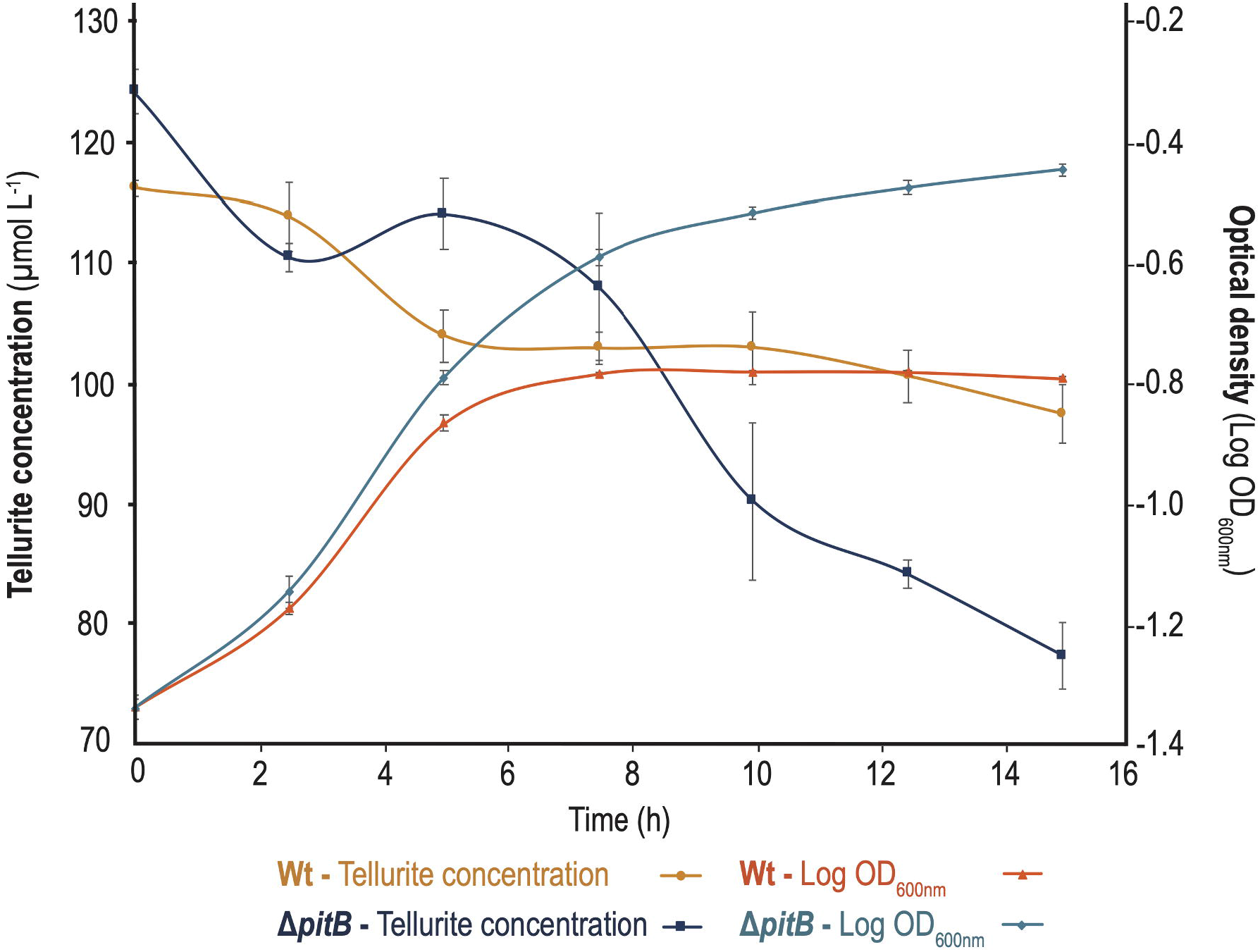
Kinetics of bacterial growth and tellurite depletion of *P. putida* KT2440 and Δ*pitB* strains in M9 minimal media supplemented with tellurite (100 μM). We determined tellurite using ICP-OES as described in the materials and methods. Three independent experiments were performed; the average is shown with its uncertainty. The strains were grown in M9 minimal media with citrate (0.2 %) as sole source of carbon and supplemented with potassium tellurite (100 μM). The Δ*pitB* strain showed greater tellurite uptake than the wild-type strain.

Our results differ from those reported for *E. coli,* for which it was observed that a Δ*pitA* mutant accumulated only half the tellurite of the wild-type strain [19]. The differences from our results are easily explained when considering the concentrations used in the experiments. For *E. coli* a MIC 50 μM was reported; tellurite uptake experiments were performed at concentration 20 μM [19]. At this concentration, the oxidative stress generated was not lethal for *E. coli*, allowing biomass formation and effective tellurite conversion. In contrast, the concentrations used in our kinetic experiment were near the lethal concentrations for *P. putida* KT2440, decreasing its ability to reduce tellurite. In bacteria, other tellurite transporters have been described, such as ActP permease in *R. capsulatus* [18,20] and in *E. coli* (21). Additional studies are required to determine the identity of the other transporters in *P. putida*, as well as the conditions under which each of them is active.

## Conclusion

The putative phosphate transporter PitB (PP1373) is involved in tellurite uptake in *P. putida* KT2440. The PitB transporter, based on its unspecific activity, was responsible for the early uptake of tellurite for the wild-type strain, subsequently leading to growth inhibition first and later to cellular death, caused by the oxidative stress produced by the metalloid. In *P. putida* KT2440, tellurium accumulates in the cytoplasm in nano-sized, rod-shaped particles. The Δ*pitB* strain is also capable of accumulating tellurium nanostructures in the cytoplasm, however, showed a remarkable resistance to this oxyanion with respect to the wild strain. Reduction kinetics measured with ICP-MS revealed that at concentration 100 μM of tellurite the wild type strain showed great sensitivity; its biomass formation decreased by half and the oxyanion reduction was only ~16 % in minimum liquid medium M9 whereas Δ*pitB* was able to grow normally and reduce ~38 % of the tellurite in the culture medium. The higher percentage of tellurite conversion of the Δ*pitB* strain is related to its greater capacity to form biomass at this concentration, instead of a greater uptake per cell. The phenotype of Δ*pitB* strain was restored on over-expressing *pitB* in trans, which validates our results. In conclusion, our results indicate that PitB is one of several transporters responsible for tellurite uptake in *P. putida* KT2440. Our work provides new evidence for tellurite metabolism in *P. putida*. Further studies of other transport systems and enzymes responsible for performing tellurium transformations are required, first to understand the reduction mechanism of the metalloid, and then to eventually develop a biotechnological system based on *P. putida* for the bioremediation of tellurite in the environment.

## Acknowledgements

We thank Victor de Lorenzo group in Centro Nacional de Biotecnología (CNB-CSIC), Madrid, Spain for providing the genetic tools for *pitB* deletion.

## Author contributions

RM, CC-R, PF, MC conceived and designed the experiments; RM, SV, DW-E, FV-C, CC-R, PF, PC, RP performed the experiments; RM, MC analyzed the data; PF, MC contributed reagents or materials or analytical tools; RM, MC wrote the paper. All authors reviewed and approved the final version of the manuscript.

## Competing financial interests

The authors declare no competing financial interests.

## Funding

The Vice-rectory of Research of Universidad de Costa Rica (project number 809-B7-A43) and Centro Nacional de Innovaciones Biotecnológicas (CENIBiot) supported this research.

## Notes

### Competing Interest Statement

The authors have declared no competing interest.

## References

1. Cucchiella F, D’Adamo I, Lenny Koh SC, Rosa P. Recycling of WEEEs: An economic assessment of present and future e-waste streams. Renew Sustain Energy Rev. 2015;51:263–272. doi: dx.doi.org/10.1016/j.rser.2015.06.010

2. Chasteen TG, Fuentes DE, Tantaleán JC, Vásquez CC. Tellurite: History, oxidative stress, and molecular mechanisms of resistance. FEMS Microbiol Rev. 2009;33(4):820–832. doi: 10.1111/j.1574-6976.2009.00177.x

3. Maltman C, Yurkov V. Extreme environments and high-level bacterial tellurite resistance. Microorganisms. 2019;7(12):1–24. doi: 10.3390/microorganisms7120601

4. Presentato A, Turner RJ, Vásquez CC, Yurkov V, Zannoni D. Tellurite-dependent blackening of bacteria emerges from the dark ages. Environ Chem. 2019;16(4):266–288. doi: doi.org/10.1071/EN18238

5. Taylor DE. Bacterial tellurite resistance. Trends Microbiol. 1999;7(3):111–115. doi: 10.1016/s0966-842x(99)01454-7

6. Maltman C, Yurkov V. Bioremediation potential of bacteria able to reduce high levels of selenium and tellurium oxyanions. Arch Microbiol. 2018;200(10):1411–1417. doi: dx.doi.org/10.1007/s00203-018-1555-6

7. Yurkov V V., Beatty JT. Aerobic Anoxygenic Phototrophic Bacteria. Microbiol Mol Biol Rev. 1998;62(3):695–724. doi: 10.1128/MMBR.62.3.695-724.1998

8. Di Tomaso G, Fedi S, Carnevali M, Manegatti M, Taddei C et al. The membrane-bound respiratory chain of *Pseudomonas pseudoalcaligenes* KF707 cells grown in the presence or absence of potassium tellurite. Microbiology. 2002;148(6):1699–1708. doi: 10.1099/00221287-148-6-1699.

9. Basnayake RST, Bius JH, Akpolat OM, Chasteen TG. Production of dimethyl telluride and elemental tellurium by bacteria amended with tellurite or tellurate. Appl Organomet Chem. 2001;15(6):499–510. doi: doi.org/10.1002/aoc.186

10. Moore MD, Kaplan S. Identification of intrinsic high-level resistance to rare-earth oxides and oxyanions in members of the class Proteobacteria: Characterization of tellurite, selenite, and rhodium sesquioxide reduction in *Rhodobacter sphaeroides*. J Bacteriol. 1992;174(5):1505–1514. doi: 10.1128/jb.174.5.1505-1514.1992

11. Forootanfar H, Amirpour-Rostami S, Jafari M, Forootanfar A, Yousefizadeh Z et al. Microbial-assisted synthesis and evaluation the cytotoxic effect of tellurium nanorods. Mater Sci Eng C. 2015;49:183–189. doi: 10.1016/j.msec.2014.12.078

12. Bao H, Lu Z, Cui X, Qiao Y, Guo J et al. Extracellular microbial synthesis of biocompatible CdTe quantum dots. Acta Biomater. 2010;6(9):3534–3541. doi: dx.doi.org/10.1016/j.actbio.2010.03.030

13. Turner RJ, Borghese R, Zannoni D. Microbial processing of tellurium as a tool in biotechnology. Biotechnol Adv. 2012;30(5):954–963. doi: dx.doi.org/10.1016/j.biotechadv.2011.08.018

14. Trutko SM, Akimenko VK, Suzina NE, Anisimova LA, Shlyapnikov MG et al. Involvement of the respiratory chain of gram-negative bacteria in the reduction of tellurite. Arch Microbiol. 2000;173(3):178–186. doi: 10.1007/s002039900123

15. Pérez JM, Calderón IL, Arenas FA, Fuentes DE, Pradenas GA et al. Bacterial toxicity of potassium tellurite: Unveiling an ancient enigma. PLoS One. 2007;2(2). doi: 10.1371/journal.pone.0000211

16. Borsetti F, Francia F, Turner RJ, Zannoni D. The thiol:disulfide oxidoreductase DsbB mediates the oxidizing effects of the toxic metalloid tellurite (TeO_3_^2−^) on the plasma membrane redox system of the facultative phototroph *Rhodobacter capsulatus*. J Bacteriol. 2007;189(3):851–859. doi: 10.1128/JB.01080-06

17. Turner R.J. Bacterial Tellurite Resistance. In: Kretsinger R.H., Uversky V.N., Permyakov E.A. (eds) Encyclopedia of Metalloproteins. Springer, New York, NY 2013. doi: doi.org/10.1007/978-1-4614-1533-6

18. Borghese R, Marchetti D, Zannoni D. The highly toxic oxyanion tellurite (TeO_3_^2−^) enters the phototrophic bacterium *Rhodobacter capsulatus* via an as yet uncharacterized monocarboxylate transport system. Arch Microbiol. 2008;189(2):93–100. doi: 10.1007/s00203-007-0297-7

19. Elías AO, Abarca MJ, Montes RA, Chasteen TG, Pérez-Donoso JM et al. Tellurite enters *Escherichia coli* mainly through the PitA phosphate transporter. Microbiology Open. 2012;1(3):259–267. doi: dx.doi.org/10.1016/j.micres.2015.04.010

20. Borghese R, Zannoni D. Acetate permease (ActP) is responsible for tellurite (TeO_3_^2−^) uptake and resistance in cells of the facultative phototroph *Rhodobacter capsulatus*. Appl Environ Microbiol. 2010;76(3):942–944. doi: 10.1128/AEM.02765-09

21. Elías A, Díaz-Vásquez W, Abarca-Lagunas MJ, Chasteen TG, Arenas F et al. The ActP acetate transporter acts prior to the PitA phosphate carrier in tellurite uptake by *Escherichia coli*. Microbiol Res. 2015;177:15–21. doi: dx.doi.org/10.1016/j.micres.2015.04.010

22. Nikel PI, Chavarría M, Danchin A, de Lorenzo V. From dirt to industrial applications: *Pseudomonas putida* as a Synthetic Biology chassis for hosting harsh biochemical reactions. Curr Opin Chem Biol. 2016;34:20–29. doi: 10.1016/j.cbpa.2016.05.011

23. Nikel PI, Martínez-García E, De Lorenzo V. Biotechnological domestication of pseudomonads using synthetic biology. Nat Rev Microbiol. 2014;12(5):368–379. doi: dx.doi.org/10.1038/nrmicro3253

24. Avendaño R, Chaves N, Fuentes P, Sánchez E, Jiménez JI et al. Production of selenium nanoparticles in *Pseudomonas putida* KT2440. Sci Rep. 2016;6:1–9. doi: http://dx.doi.org/10.1038/srep37155

25. Belda E, van Heck RGA, José López-Sánchez M, Cruveiller S, Barbe V et al. The revisited genome of *Pseudomonas putida* KT2440 enlightens its value as a robust metabolic chassis. Environ Microbiol. 2016;18(10):3403–3424. doi: 10.1111/1462-2920.13230

26. Nikel P, Chavarría M. Quantitative Physiology Approaches to Understand and Optimize Reducing Power Availability in Environmental Bacteria. In: Hydrocarbon and Lipid Microbiology Protocols - Springer Protocols Handbooks. Springer P. Springer, Berlin, Heidelberg; 2015. p. 39–70. doi: doi.org/10.1007/8623_2015_84

27. Nikel PI, Chavarría M, Fuhrer T, Sauer U, De Lorenzo V. *Pseudomonas putida* KT2440 strain metabolizes glucose through a cycle formed by enzymes of the Entner-Doudoroff, embden-meyerhof-parnas, and pentose phosphate pathways. J Biol Chem. 2015;290(43):25920–32. doi: 10.1074/jbc.M115.687749

28. Nikel PI, Fuhrer T, Chavarría M, Sanchez-Pascuala A, Sauer U et al. Redox stress reshapes carbon fluxes of *Pseudomonas putida* for cytosolic glucose oxidation and NADPH generation. bioRxiv. 2020;2020.06.13.149542. doi: https://doi.org/10.1101/2020.06.13.149542

29. Sánchez-Romero JM, Díaz-Orejas R, De Lorenzo V. Resistance to tellurite as a selection marker for genetic manipulations of Pseudomonas strains. Appl Environ Microbiol. 1998;64(10):4040–4046. doi: 10.1128/AEM.64.10.4040-4046.1998

30. Rajwade JM, Paknikar KM. Bioreduction of tellurite to elemental tellurium by *Pseudomonas mendocina* MCM B-180 and its practical application. Hydrometallurgy. 2003;71(1-2):243–248. doi: https://doi.org/10.1016/S0304-386X(03)00162-2

31. Aparicio T, de Lorenzo V, Martínez-García E. CRISPR/Cas9-Based counterselection boosts recombineering efficiency in *Pseudomonas putida*. Biotechnol J. 2018;13(5):1–10. doi: 10.1002/biot.201700161

32. Sambrook J, Russell D. Molecular Cloning: a laboratory manual. 3rd ed. Cold Spring Harbor; 2001. Isbn: 978-087969577-4

33. Silva-Rocha R, Martínez-García E, Calles B, Chavarría M, Arce-Rodríguez A et al. The Standard European Vector Architecture (SEVA): A coherent platform for the analysis and deployment of complex prokaryotic phenotypes. Nucleic Acids Res. 2013;41(D1):666–675. doi: doi.org/10.1093/nar/gks1119

34. Jackson RJ, Binet MRB, Lee LJ, Ma R, Graham AI et al. Expression of the PitA phosphate/metal transporter of *Escherichia coli* is responsive to zinc and inorganic phosphate levels. FEMS Microbiol Lett. 2008;289(2):219–224. doi: 10.1111/j.1574-6968.2008.01386.x

35. Turner RJ, Weiner JH, Taylor DE. Tellurite-mediated thiol oxidation in *Escherichia coli*. Microbiology. 1999;145(9):2549–2557. doi: 10.1099/00221287-145-9-2549

36. Turner RJ, Aharonowitz Y, Weiner JH, Taylor DE. Glutathione is a target in tellurite toxicity and is protected by tellurite resistance determinants in *Escherichia coli*. Can J Microbiol. 2001;47(1):33–40. doi: 10.1139/w00-125

37. Chavarría M, Nikel PI, Pérez-Pantoja D, De Lorenzo V. The Entner-Doudoroff pathway empowers *Pseudomonas putida* KT2440 with a high tolerance to oxidative stress. Environ Microbiol. 2013;15(6):1772–1785. doi: 10.1111/1462-2920.12069

38. Presentato A, Piacenza E, Anikovskiy M, Cappelletti M, Zannoni D et al. *Rhodococcus aetherivorans* BCP1 as cell factory for the production of intracellular tellurium nanorods under aerobic conditions. Microb Cell Fact. 2016;15(1):1–14. doi: 10.1186/s12934-016-0602-8

