## Supplementary information for "The putative phosphate transporter PitB (PP1373) is involved in tellurite uptake in *Pseudomonas putida* KT2440"

By

Rafael Montenegro[1*], Sofía Vieto[1*], Daniela Wicki-Emmenegger[1,2,3], Felipe Vásquez-Castro[1,2], Carolina Coronado-Ruiz[1,2], Paola Fuentes-Schweizer[2,4] Paula Calderón[5], Reinaldo Pereira[6], & Max Chavarría[1,2,3]

[1]Centro Nacional de Innovaciones Biotecnológicas (CENIBiot), CeNAT-CONARE, 1174-1200 San José, Costa Rica. [2]Escuela de Química, Universidad de Costa Rica, 11501-2060 San José, Costa Rica. [3]Centro de Investigaciones en Productos Naturales (CIPRONA), Universidad de Costa Rica, 11501-2060 San José Costa Rica. [4]Centro de Electroquímica y Energía Química (CELEQ), Universidad de Costa Rica, 11501-2060 San José, Costa Rica. [5]Centro de Investigaciones en Estructuras Microscópicas (CIEMIC), Universidad de Costa Rica, 11501-2060 San José, Costa Rica. [6]Laboratorio Nacional de Nanotecnología (LANOTEC), CeNAT-CONARE, 1174-1200 San José, Costa Rica.

***Keywords:*** *pitB*, phosphate transporter, tellurite, *Pseudomonas putida*, tellurium nanoparticles.

---

* These authors contributed equally to this work.

* Correspondence to: Max Chavarría

Escuela de Química & Centro de Investigaciones en Productos Naturales (CIPRONA)

Universidad de Costa Rica

Sede Central, San Pedro de Montes de Oca

San José, 11501-2060, Costa Rica

Phone (+506) 2511 8520. Fax (+506) 2253 5020

ORCID: https://orcid.org/0000-0001-5901-3576

### Supplementary Materials and Methods

#### Pit protein homology analysis

Protein sequences were retrieved from the UNIPROT database while the PitB sequence from *P. putida* was obtained from the *Pseudomonas* genome database[1]. The PRALINE software was used setting a BLOSUM62 scoring matrix[2] and having as input the desired Pit protein sequences. After the run, the PRALINE software generated the alignment and the identity scores to finally do the homology analysis. Initially, the analysis was performed to compare the Pit proteins between *E. coli* and *P. putida*, following, the same analysis was performed to compared Pit proteins among different *Pseudomonas* species.

#### Phylogenetic analysis based on the Pit proteins

Protein sequences were retrieved from the UNIPROT database while the PitB from *P. putida* was obtained from the *Pseudomonas* genome database[1]. Then, the Clustal Omega tool[3] was used to create a multiple alignment of the Pit protein sequences. Finally, the phylogenetic tree was created by a maximum-likelihood method based on the general time-reversible model using MEGA software[4]. The multiple alignment file previously made was the input and the bootstrap replications were set to be 100 in total, values that are represented in the phylogenetic tree.

### Legends of Supplementary Figures

**Figure S1. Alignment of Pit protein sequences in order to identify homologies between *E. coli* and *P. putida* proteins**. The PRALINE software was used setting a BLOSUM62 scoring matrix to generate the alignment and the homology analysis. Pit proteins from *E. coli* share a high identity between them, nevertheless they both have lower identities compared to the PitB from *P. putida*, showing no affinity for anyone in particular.

**Figure S2. Alignment of Pit protein sequences from different *Pseudomonas* species.** The PRALINE software was used to generate the alignment and the homology analysis setting a BLOSUM62 scoring matrix.

Analysis that revealed a high identity between the Pit proteins from *P. aeruginosa*, *P. syringae* and *P. fluorescens* compared to the PitB protein from *P. putida*.

**Figure S3. Phylogenetic classification of the Pit protein in several bacterial strains focusing in *P. putida* PitB protein**. The evolutionary history was inferred from the multiple alignment using a maximum-likelihood method, using the MEGA software. Bootstrap values, 100 bootstrap replications in total, are represented in the phylogenetic tree. The phylogeny showed a higher evolutionary relationship of the PitB protein from *P. putida* to the PitA proteins from the different *Pseudomonas* species rather than to the *E. coli* PitB protein.

**Figure S4. Growth curves of *P. putida* KT2440 wild type and ∆*pitB* strains in M9 minimal media with tellurite at different concentrations**. Growth curves in different concentrations of potassium tellurite from 0 µM to 1000 µM for the wild type **(A)** and the ∆*pitB* **(B)** strains, results obtained from a 96 wells-microplate experiment over three replicates. The phenotypical analysis showed a progressively growth inhibition for the wild type strain starting at 75 µM of tellurite, while the ∆*pitB* exhibited the same inhibition, nevertheless, starting at a concentration of 200 µM.

### Supplementary References


Winsor, G. L. et al. Enhanced annotations and features for comparing thousands of Pseudomonasgenomes in the Pseudomonas genome database. Nucleic Acids Res. 44, D646–D653 (2016).

Simossis, V. A., Kleinjung, J. & Heringa, J. Homology-extended sequence alignment. Nucleic Acids Res. 33, 816–824 (2005).

Madeira, F. et al. The EMBL-EBI search and sequence analysis tools APIs in 2019. Nucleic Acids Res. 47, W636–W641 (2019).

Kumar, S., Stecher, G., Li, M., Knyaz, C. & Tamura, K. MEGA X: Molecular evolutionary genetics analysis across computing platforms. Mol. Biol. Evol. 35, 1547–1549 (2018).
