## Supplementary figures and images for "The putative phosphate transporter PitB (PP1373) is involved in tellurite uptake in *Pseudomonas putida* KT2440"

### Supp. Figure S1

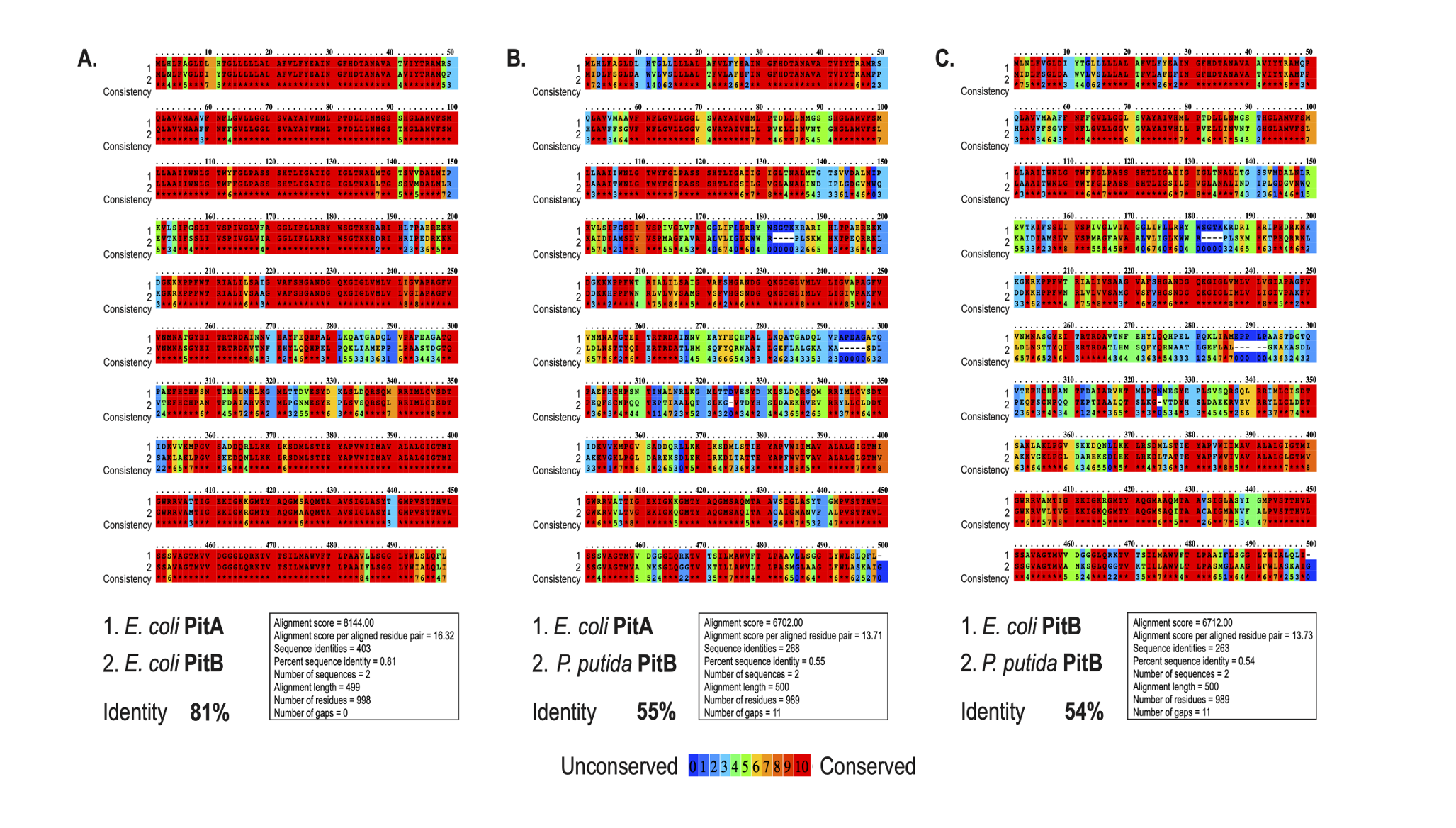

### Supp. Figure S2

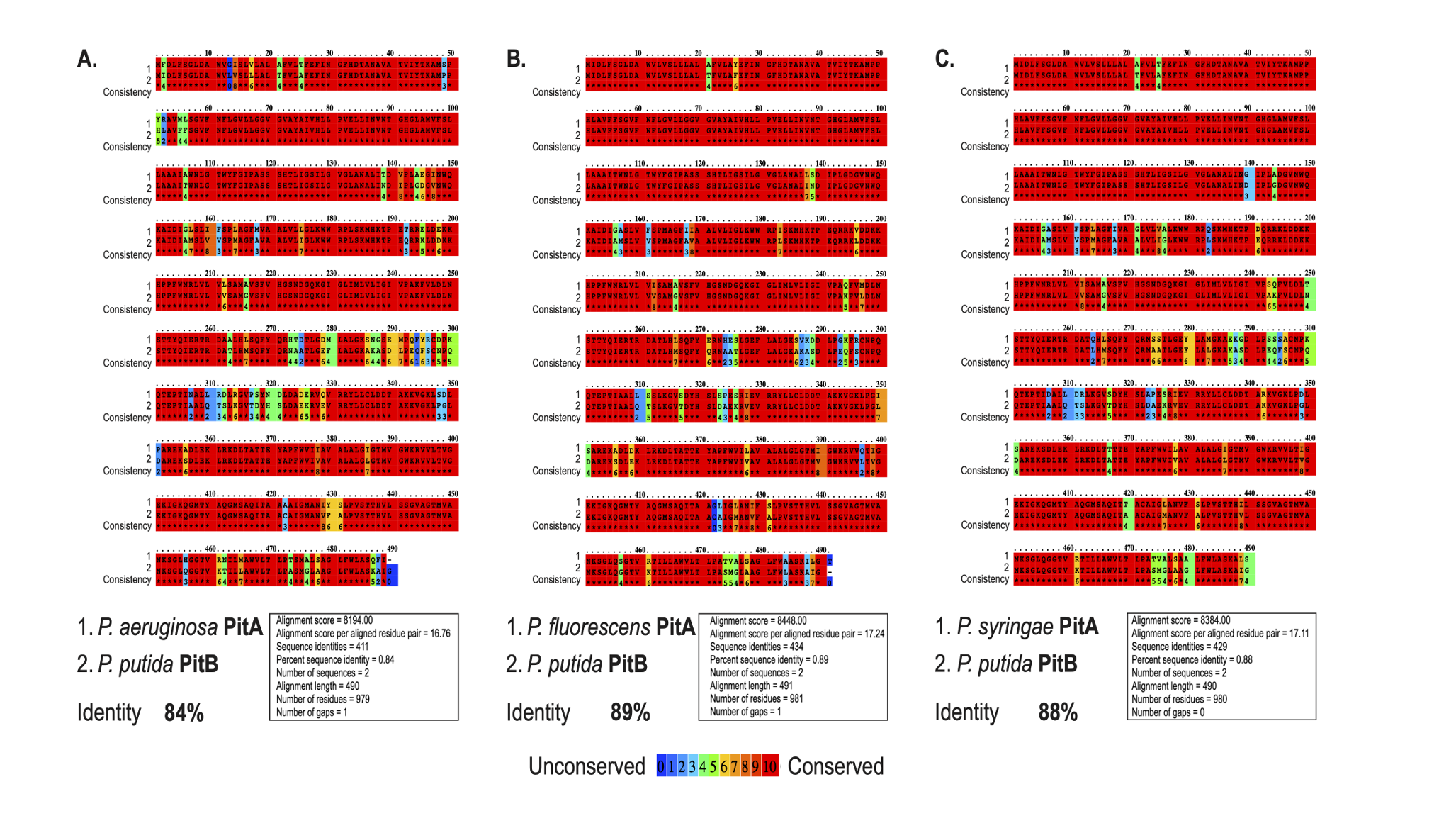

### Supp. Figure S3

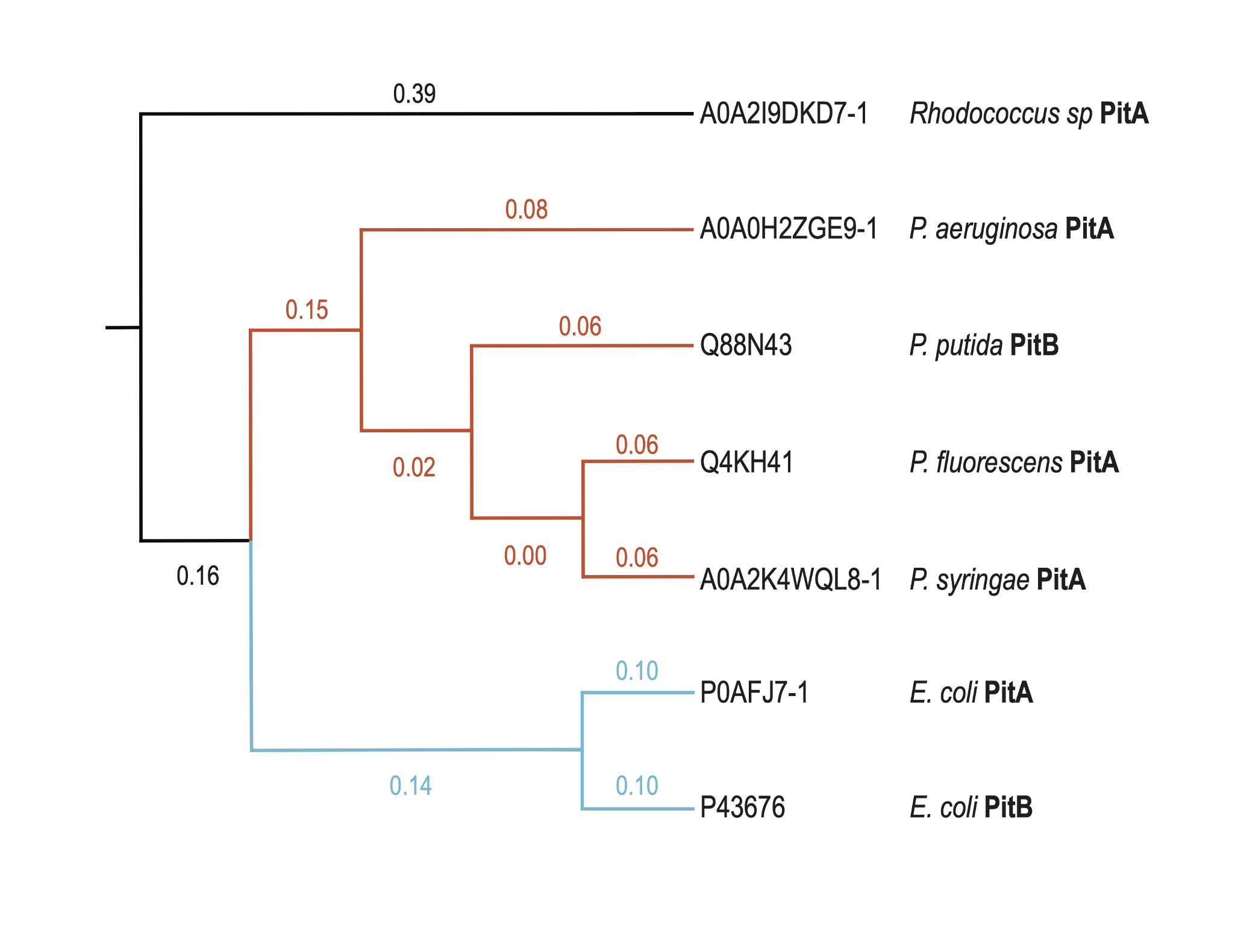

### Supp. Figure S4

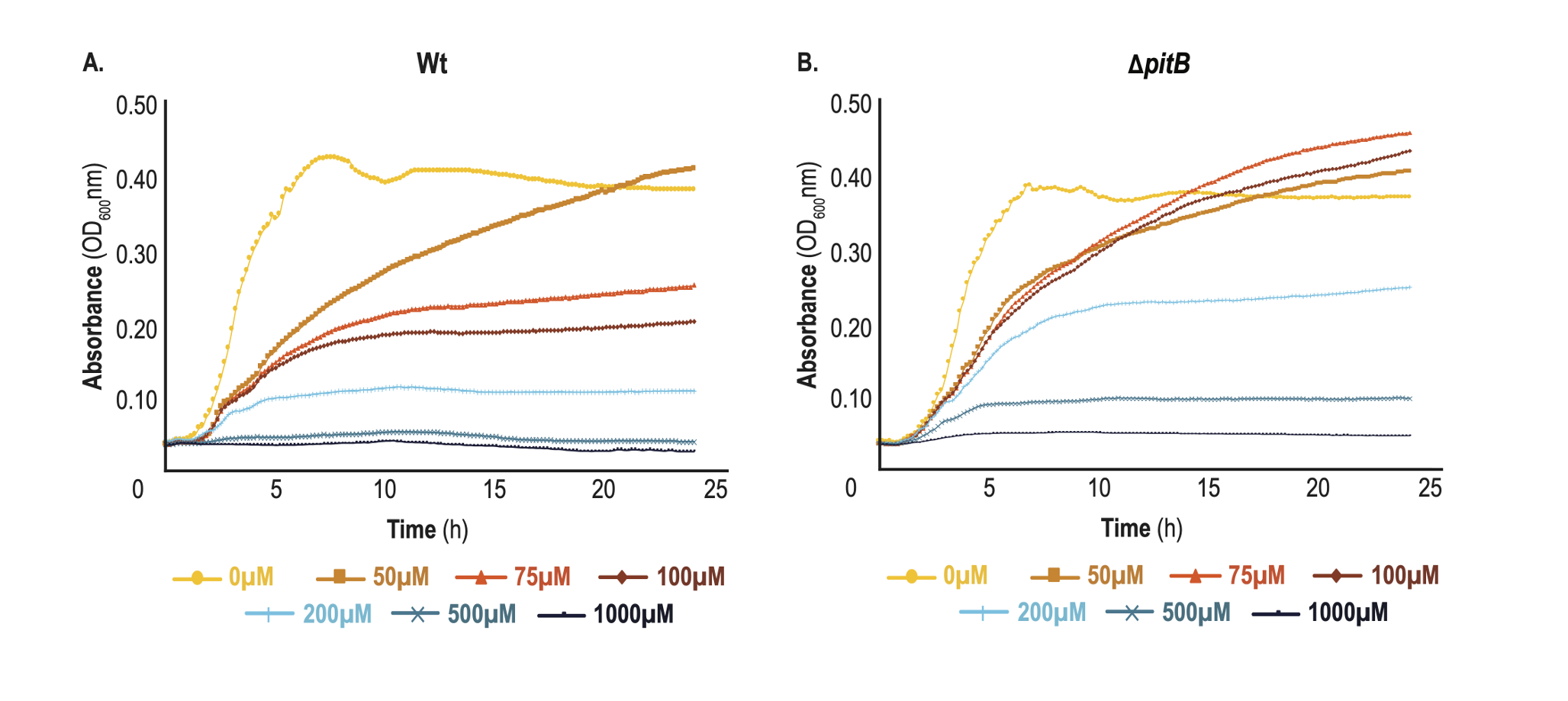
